# SHP2 Inhibition Abrogates MEK inhibitor Resistance in Multiple Cancer Models

**DOI:** 10.1101/307876

**Authors:** Carmine Fedele, Hao Ran, Brian Diskin, Wei Wei, Jayu Jen, Kiyomi Araki, Diane M Simeone, George Miller, Benjamin G Neel, Kwan Ho Tang

## Abstract

Adaptive resistance to MEK inhibitors (MEK-Is) typically occurs via induction of genes for different receptor tyrosine kinases (RTKs) and/or their ligands, even in tumors of the same histotype, making combination strategies challenging. SHP2 (*PTPN11*) is required for RAS/ERK pathway activation by most RTKs, and might provide a common resistance node. We found that combining the SHP2 inhibitor SHP099 with a MEK-I inhibits proliferation of multiple cancer cells *in vitro*. *PTPN11* knockdown/MEK-I had similar effects, while expressing SHP099-binding mutants conferred resistance, demonstrating that SHP099 was on-target. This combination was efficacious in xenograft and/or genetically engineered models of *KRAS*-mutant pancreas cancer and ovarian cancer and in wild-type RAS-expressing triple negative breast cancer. Biochemical studies show that SHP099 impedes SOS/RAS/MEK/ERK1/2 reactivation in response to MEK-Is and blocks ERK1/2-dependent transcriptional programs. SHP099 alone also inhibited RAS activation in some, but not all, *KRAS*-mutant lines. Hence, SHP099/MEK-I combinations could have therapeutic utility in multiple malignancies.

**SIGNIFICANCE:** MEK inhibitors have shown limited efficacy as single agents because of the rapid development of adaptive resistance. We find that combining SHP2 and MEK inhibition abrogates adaptive resistance in multiple cancer models, expressing mutant and wild-type KRAS.

## INTRODUCTION

The RAS/ERK mitogen-activated protein kinase (MAPK) pathway is one of the most commonly affected signaling pathways in human cancer (1-3). Mutations in genes encoding pathway components, including those for receptor tyrosine kinases (RTKs), SHP2, NF1, RAS or RAF, cause inappropriate pathway activation and promote oncogenesis. Attempts have been made to target the ERK pathway in different cancer types, and can lead to initial responses. Unfortunately, “adaptive resistance” occurs in most patients, leading to recurrence or progression (4).

*KRAS* is the most frequently mutated RAS/ERK pathway gene (1-3). Approaches to target *KRAS*-mutant cancers with MEK-inhibitors (MEK-Is) have failed, often due to the induction of RTK genes and/or their ligands. For example, FGFR1 is activated in MEK-I-treated *KRAS*-mutant lung cancers, leading to increased upstream signaling and ERK reactivation (5). Another group found that MEK-I resistance can be mediated through ERBB3 in *KRAS*-mutant lung and colon cancers (6), whereas a third reported that MEK-I treatment leads to EGFR activation in *KRAS* mutant pancreatic cancer lines (7). Malignancies that lack mutations in pathway genes but nonetheless hyperactivate ERK also show adaptive resistance in response to MEK-Is. For example, MEK-I-treated triple negative breast cancer (TNBC) cells induce the expression of genes encoding AXL, DDR1, FGFR2, IGF1R, KIT, PDGFRB and VEGFRB (8,9).

Because resistance to MEK-Is can be mediated by a variety of RTKs, combining MEK and RTK inhibition is not likely to be a viable therapeutic approach. Instead, a strategy that efficiently blocks signals from multiple activated RTKs is required to abolish adaptive resistance to MEK-Is. The protein-tyrosine phosphatase SHP2 is a positive (i.e., signal-enhancing) signal transducer, acting between RTKs and RAS (10,11). A potent, highly specific inhibitor targeting SHP2, SHP099, has been developed, and blocks ERK activation and proliferation of cancer cells driven by over-expressed, hyperactivated RTKs (12,13). We hypothesized that SHP099 would inhibit signals from RTKs activated following MEK inhibition, and thereby block adaptive resistance. This idea comports with the previous finding that *PTPN11* shRNA or CRISPR/Cas9-mediated deletion prevents adaptive resistance to vemurafenib in *BRAF*-mutant colon cancer (14).

Here, we test this hypothesis in multiple *KRAS*-mutant and wild type cancer cells from different histotypes. Our results suggest that SHP2 inhibition could provide a general strategy for preventing MEK-I resistance in a wide range of malignancies.

## RESULTS

### SHP099 abrogates adaptive resistance to MEK-inhibitors *in vitro*

Previous work showed that various cancer models develop adaptive resistance to MEK-Is through RTK upregulation. We analyzed RTKs/RTK ligand gene expression by qRT-PCR in pancreatic ductal adenocarcinoma (PDAC) cell lines treated with AZD6244 (Fig. 1A and B). Consistent with the earlier findings, several—but different–RTKs were induced by MEK-I treatment, including *EGFR*, *FGFR3*, *IGFR1* and *MET* in MIAPaCa-2 cells, *ERBB2/3*, *FGFR1/2* and *IGFR1* in Capan-2 cells, and *ERBB2/3* and *FGFR3* in CFPAC-1 cells (Fig. 1A). The same lines variably induced *EGF*, *FGF2, PDGFB*, *PDGFD* and/or *VEGFA/B* RNA (Fig. 1B). These observations make it difficult, if not impossible, to design an efficient combination therapy with MEK-Is by targeting RTKs directly.

**Figure 1.**
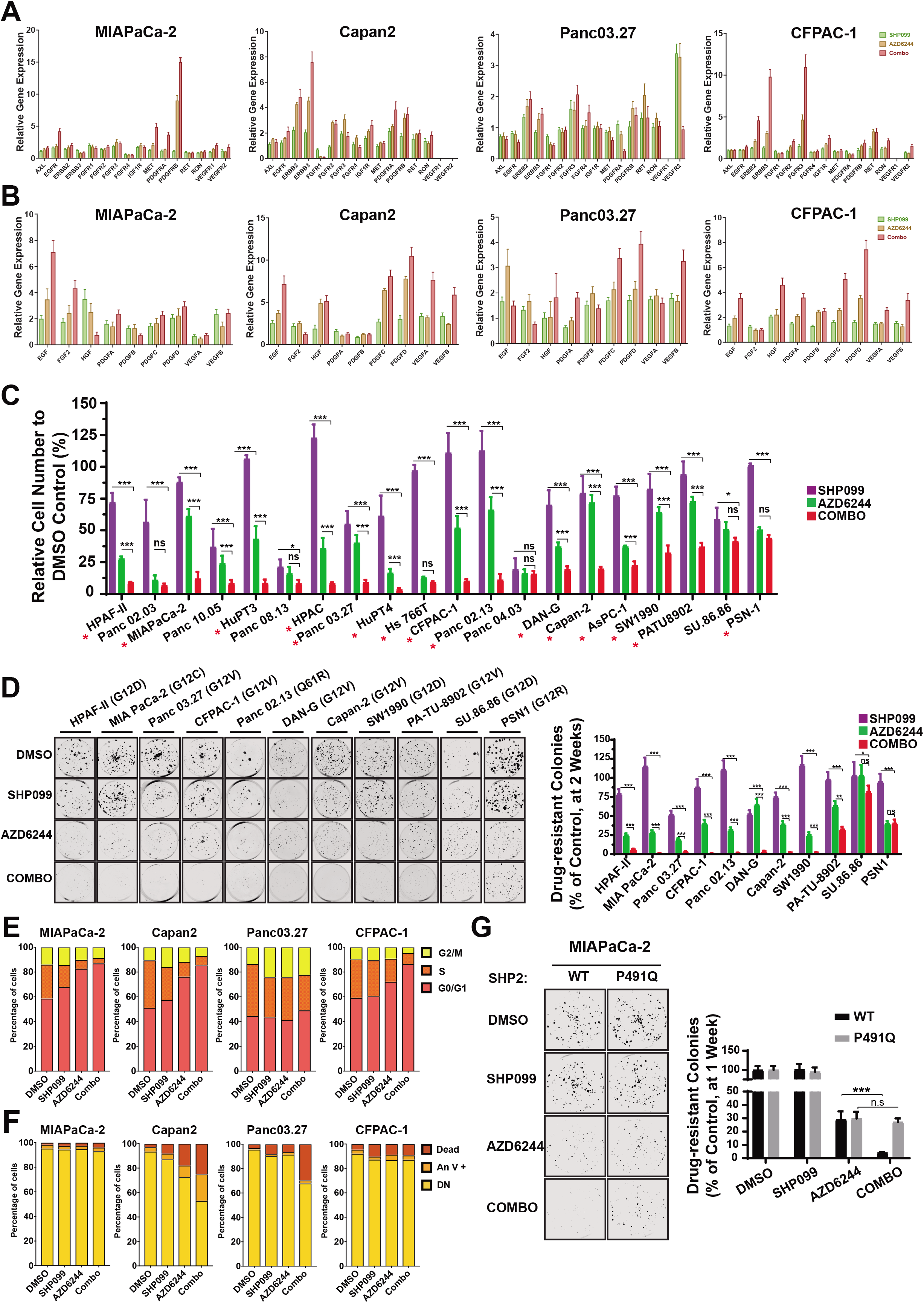
Combined SHP2 and MEK inhibition abrogates adaptive resistance in PDAC cell lines. **A-B**, Time-dependent increase in in RTK (A) and RTK ligand (B) gene expression in PDAC cells after DMSO, SHP099, AZD6244, or SHP099/AZD6244 (Combo) treatment, determined by qRT-PCR. **CD**, PDAC cell models were treated with DMSO, SHP099, AZD6244, or both drugs (Combo). Cell viability by PrestoBlue assay (C) and colony formation (D) were assessed at one or two weeks, respectively (\**P*< 0.05, \*\**P*< 0.001, \*\*\**P*< 0.0001, two-sided *t* test). Representative results from a minimum of three biological replicates are shown per condition. Red asterisks indicate synergistic interaction between the two drugs by BLISS independent analysis. **E-F**, Cell cycle (E) and cell death analysis (F) of cells treated as in C by flow cytometry with Annexin V/PI staining, following 48 hours of drug treatment. **G**, Colony formation assay (one week) in MiaPaCa-2 cells either expressing an SHP099-resistant *PTPN11* mutant (P491Q) or wild-type *PTPN11* (WT) (\*\**P*<0.0001, two-sided *t* test). Representative results from a minimum of three biological replicates are shown per condition. For all experiments, drug doses were: SHP099 10 μM, AZD6244 1 μM, Combo= SHP099 10 μM + AZD6244 1μM. Numbers under blots indicate relative intensities, compared with untreated controls, quantified by LICOR.

To explore whether SHP2 inhibition could suppress MEK-I adaptive resistance, we first performed *in vitro* viability (PrestoBlue) and colony formation assays on a panel of *KRAS*-mutant PDAC lines (Fig. 1C and D). Resistant cell populations and drug-resistant colonies were observed after one or two weeks, respectively, of treatment with a well-established MEK-I, AZD6244. AZD6244 itself had variable effects, leading to 30-90% reduction in proliferation/colony formation compared with control DMSO treatment; nevertheless, nearly all lines showed significant resistance after AZD6244 treatment (Fig. 1C and D). Consistent with the conclusions of a previous report (12), these *KRAS*-mutant cell lines exhibited low sensitivity to SHP099 alone. By contrast, all but two of the lines had markedly reduced cell numbers and few or no detectable colonies after MEK-I/SHP099 combination treatment, and in most cases, the combination showed synergistic efficacy (Fig. 1C, red asterisks). Similar effects were seen in full growth curve assays (Fig. S1A) or when a different MEK-I (trametinib) was used (data not shown). The drug combination decreased cell cycle progression and, in some lines, enhanced cell death (measured at 48h and 6 days of treatment respectively), compared with either single agent alone (Fig. 1E and F). Similar inhibitory effects of the combination were seen in a small sample of pancreas cancer cells from patient-derived xenografts (PDXs) in short-term cultures and in the *KRAS*-mutant non-small cell lung cancer (NSCLC) lines that we tested (Fig S1B-D).

To test whether the effects of SHP099 reflected SHP2 inhibition and not an off-target effect, we expressed a *PTPN11* mutant (*PTPN11*^*P491Q*^) predicted to lack drug binding in MiaPaCa-2 cells, which are quite sensitive to the SHP099/MEK-I. Reassuringly, expression of this mutant eliminated the effects of SHP099 in MEK-I/SHP099-treated cells (Fig. 1G). Another drug-resistant mutant, *PTPN11*^*T253M/Q257L*^, rescued the effects of the combination on H358 NSCLC cells (Fig. S1E). Moreover, combining MEK inhibition with expression of a *PTPN11* shRNA had similar effects as MEK-I/SHP099 treatment (Fig. S1F). Taken together, these data suggest that SHP099 is “on target,” and that SHP2 inhibition diminishes adaptive resistance to MEK-Is in multiple *KRAS*-mutant cancer cell lines, arising from two distinct tissues.

### SHP099 impedes MEK inhibitor-induced reactivation of the ERK MAPK pathway

We next assessed the biochemical effects of each single agent and the drug combination on RAS/ERK pathway activation after short (1h)- and longer-term (48h) treatment. As expected from previous reports (Introduction) and the induction of multiple RTKs/RTK ligands seen above, short-term AZD6244 treatment had no detectable effect on RAS. After 48 h, however, RAS activation (as monitored by RAF-RBD pull down assay) was enhanced (Fig. 2A). Analysis using isoform-specific RAS antibodies revealed increased activation of KRAS and NRAS in response to 48h MEK-I treatment of MiaPaCa-2 cells (Fig. 2B). These cells express *KRAS*^*G12C*^, but not WT *KRAS* (mutant allele frequency=0.99) (15,16), so the increase seen in KRAS activation must reflect enhanced cycling of the mutant KRAS. KRAS(G12C) retains significant intrinsic GTPase activity (17), and our findings indicate that this activity contributes significantly to the steady state level of mutant-KRAS-GTP in these cells. By contrast, the increase in NRAS-GTP in response to inhibitor treatment reflects activation of normal, endogenous NRAS.

**Figure 2.**
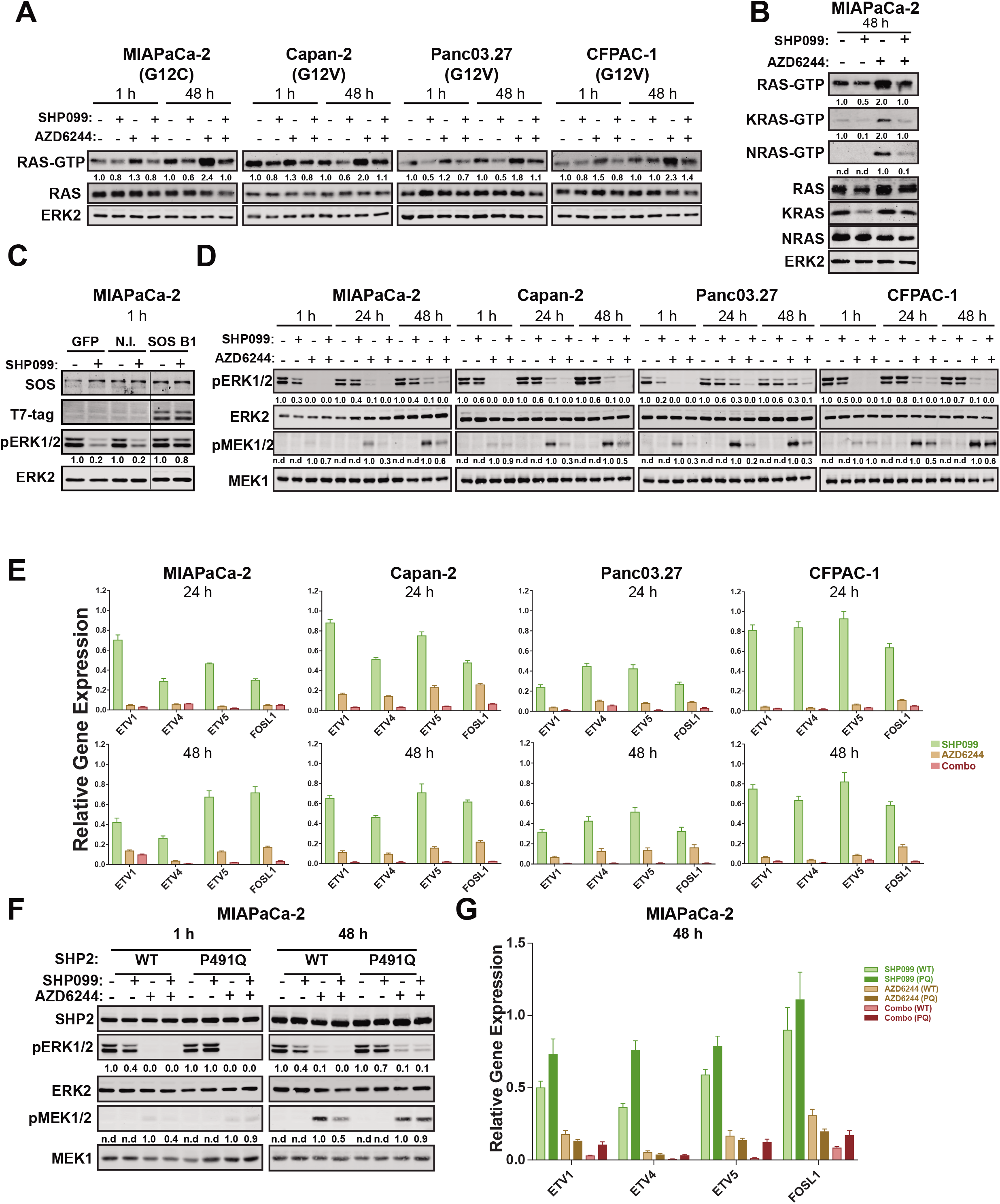
SHP2 inhibition acts upstream of RAS to abrogate MEK-I-evoked ERK MAPK pathway reactivation. **A-B**, Immunoblots of whole cell lysates or GST-RBD-precipitated (RAS-GTP, KRASGTP and NRAS-GTP) lysates from PDAC cells treated with DMSO, SHP099 10 μM, AZD6244 1 μM, or both drugs for the times indicated. The images shown are representative of at least two independent biological replicates. **C**, Effect of SHP099 on p-ERK levels in MiaPaCa-2 cells expressing a SOS1 mutant (SOS B1) that targets the SOS1 catalytic domain constitutively to the plasma membrane. Cells were incubated for 1 hour with SHP099, and lysates wre immunoblotted for p-ERK and total ERK (as a loading control). **D**, Immunoblots of lysates from PDAC lines treated as indicated. Image shown is representative of three independent biological replicates. **E**, ERK-dependent gene expression levels (*ETV1,4, 5* and *FOSL-1*), assessed by qRT-PCR, in PDAC that for the indicated timess. **F**, Immunoblot analysis of SHP2, p-ERK, ERK, p-MEK and MEK from MiaPaCa-2 cells ectopically-expressing wild-type SHP2 (WT) or an SHP099 resistant-mutant (P491Q), treated with the indicated drugs and times. **G**, ERK-dependent gene expression in MiaPaCa-2 cells ectopically expressing wild-type SHP2 (WT) or an SHP099-resistant mutant (P491Q), treated as described in F. Numbers under blots indicate relative intensities, compared with untreated controls, quantified by LICOR.

SHP2 is required for RTK-evoked RAS activation, and consequently, for MEK and ERK activation (10,11). Accordingly, SHP099 suppressed the increased levels of RAS-GTP in response to 48h MEK-I treatment in multiple *KRAS*-mutant PDAC lines (Fig. 2A). Isoform-specific immunoblots revealed that SHP099 blocked reactivation of mutant KRAS(G12C) and endogenous NRAS (Fig. 2B) in MiaPaCa-2 cells. The other cell lines tested express KRAS mutants that have less intrinsic GTPase activity than KRAS(G12C) (17), and the allele fraction of mutant *KRAS* in these lines is less than one. Hence, it is not clear whether SHP099 blocks activation of these RAS mutants as well, or only affects the remaining endogenous KRAS or other RAS isoforms (Fig. 2A). Single agent SHP099 also variably decreased RAS-GTP levels at 1h and 48h of treatment (Fig. 2A and B).

Although SHP2 acts upstream of RAS, whether it acts by promoting RAS exchange (e.g., via SOS) or inhibits RAS-GAP action, or even both, is less clear (10,11). As KRAS(G12C) is highly resistant to RAS-GAP (17), the decrease in KRAS-GTP seen in MiaPaCa-2 cells treated with SHP099 (+AZD6244) suggested that SHP099 acts upstream of SOS to promote RAS activation. Consistent with this notion, expressing the SOS1 catalytic domain with a C-terminal CAAX tag (18) rescued the effects of SHP099 in SHP099-treated cells (Fig. 2C). These data show unambiguously that SHP2 acts upstream of SOS in MiaPaCa-2 cells, although they do not rule out the possibility of additional effects on RAS-GAP in the context of normal RAS (see Discussion).

Consistent with the above observations, single agent AZD6244 blocked MEK and ERK1/2 phosphorylation after 1h, but these effects were successively abolished after 24h and 48h of treatment, respectively, and MEK and ERK activity rebounded. This adaptive increase in MEK and ERK phosphorylation was abolished by co-administration of SHP099 (Fig. 2D and S2A). Consistent with its single agent effects on RAS, SHP099 alone had variable effects on MEK-ERK activation. We also asked whether these biochemical events translated into effects on ERK1/2-dependent gene expression by quantifying FOS-like 1 (*FOSL1)* and ETS variants 1, 4-5 (*ETV1, 4 and 5*) RNA by qRT-PCR; ERK-dependent gene expression can also provide a more sensitive assessment of ERK pathway output than quantifying p-ERK levels (19). Compared with the effects of either single agent, the SHP099/AZD6244 combination caused greater suppression of ERK-dependent transcription (Fig. 2E). Together, these findings confirm that ERK reactivation is a key component of the survival program activated in *KRAS*-mutant cancer cells treated with MEK-Is, and suggest that SHP099 effectively suppresses this adaptive response.

Importantly, the effects of SHP099 (like its effects on colony formation; Fig. 1G) were reversed in MiaPaCa-2 cells expressing the drug-resistant mutant *PTPN11*^*P491Q*^ (Fig. 2F and G) and in *KRAS*-mutant H358 NSCLC cells expressing *PTPN11*^*T253M/Q257L*^ (Fig. S2B). Furthermore, SHP2 depletion had similar biochemical effects as SHP2 inhibition in MiaPaCa-2 cells (Fig. S2C). Hence, the synergistic effects of SHP099/MEK-I reflect on-target inhibition of SHP2.

We also explored the biochemical mechanism for resistance of two *KRAS*-mutant PDAC lines to the MEK-I/SHP099 combination. One line (PSN1), failed to suppress MEK-ERK reactivation in response to the combination (Fig S3A), and accordingly, failed to suppress ERK-dependent gene expression (Fig. S3B). By contrast, another resistant line, SU.86.86, suppressed MEK/ERK and ERK-dependent genes to an extent similar to sensitive cells, consistent with a downstream escape mechanism (Fig. S3C and D). Further investigation will be required to uncover the precise molecular explanation for resistance in these cell lines.

### Combined SHP2/MEK inhibition suppresses *KRAS*-mutant tumor growth *in vivo*

To ask whether these *in vitro* results translate to *in vivo* efficacy, we established MiaPaCa-2 and Capan-2 xenografts, and treated them with vehicle control (methyl-cellulose+Tween80), the MEK-inhibitor trametinib alone, SHP099 alone, or a trametinib/SHP099 combination. We used trametinib for these studies because of its favorable pharmacokinetic properties in mice (t_½_ = 33h (20)), which, like SHP099 (13), enables single daily dosing. In initial experiments, mice were treated daily with trametinib (1mg/kg), a dose used commonly in mouse tumor studies (6,21,22), SHP099 (75mg/kg), or both drugs. Each single agent was well-tolerated, but mice receiving the combination lost weight (>10%), exhibited lassitude, and began dying at day 7 of treatment. Some mice showed frank evidence of GI bleeding (Fig. S4A). Histological analysis of sick mice revealed multi-focal ulceration in the GI tract, acute esophagitis and gastritis, which would likely result in bleeding (Fig. S4B). Blunting of villi was also observed in the small intestine of some treated mice, which could cause malabsorption, diarrhea and weight loss (Fig. S4B).

Although trametinib is frequently administered to mice at this dose, or even at doses as high as 3mg/kg (5,23), by allometric scaling, the mouse equivalent of the maximum tolerated dose (MTD) in humans is actually ~0.25mg/kg (24). We treated a small group of mice with this dose of trametinib and SHP099 (75mg/kg) daily, but although treated mice lived longer than with the higher trametinib dose, this combination also led to weight loss and death (data not shown). After further exploratory dose-finding, we arrived at a tolerable schedule, in which trametinib was delivered at 0.25mg/kg, while SHP099 was given at its full dose (75mg/kg), but every other day (QOD). A few mice treated with this regimen developed mild, self-limited, non-bloody diarrhea, but all showed stable weight, normal behavior and appeared healthy for up to 40 days of continuous treatment (Fig. S4C and D, and data not shown).

We allowed MiaPaCa-2 (Fig. 3A) or Capan2 (Fig. 3B) xenografts to grow to 500mm^3^, and then began treatment with vehicle, trametinib (0.25mg/kg QD), SHP099 (75 mg/kg QD), or trametinib 0.25mg/kg QD, SHP099 75mg/kg QOD. Remarkably, the combination caused substantial regressions in all treated mice bearing either xenograft (Fig. 3A and B), with tumor shrinkage averaging >65% in both xenograft models, well above Response Evaluation Criteria for Solid Tumors (RECIST) criteria (25). The effects of the single agents were more variable both within each treatment group and in mice bearing MiaPaCa-2 versus Capan2-derived tumors. Trametinib had minimal effects on MIAPaCa-2 tumors (Fig. 3A), whereas it caused significant shrinkage of about half of the Capan2 xenografts (Fig. 3B). Only a few trametinib-treated mice met RECIST criteria (>30%), however, and the SHP099/MEK-I combination was clearly more effective (Fig. 3A and B). Surprisingly, in contrast to its lack of effect on proliferation in cell culture or on colony formation, SHP099 alone caused tumor shrinkage in ~60% of MIAPaCa-2 and ~80% of Capan2 xenografts. It is not clear whether this discrepancy stems from effects of SHP099 on normal RAS in cells within the tumor microenvironment (e.g., fibroblasts, blood vessels) or reflects SHP099 effects on the tumor cells themselves (see Discussion). Nevertheless, single agent SHP099 was, like trametinib alone, clearly inferior to the drug combination. Consistent with these biological effects, immunoblotting (Fig. 3C) and immunohistochemical (IHC) analysis (Fig. 3D) revealed more profound p-ERK inhibition in combination-treated tumors than in those receiving trametinib or SHP099 alone.

**Figure 3.**
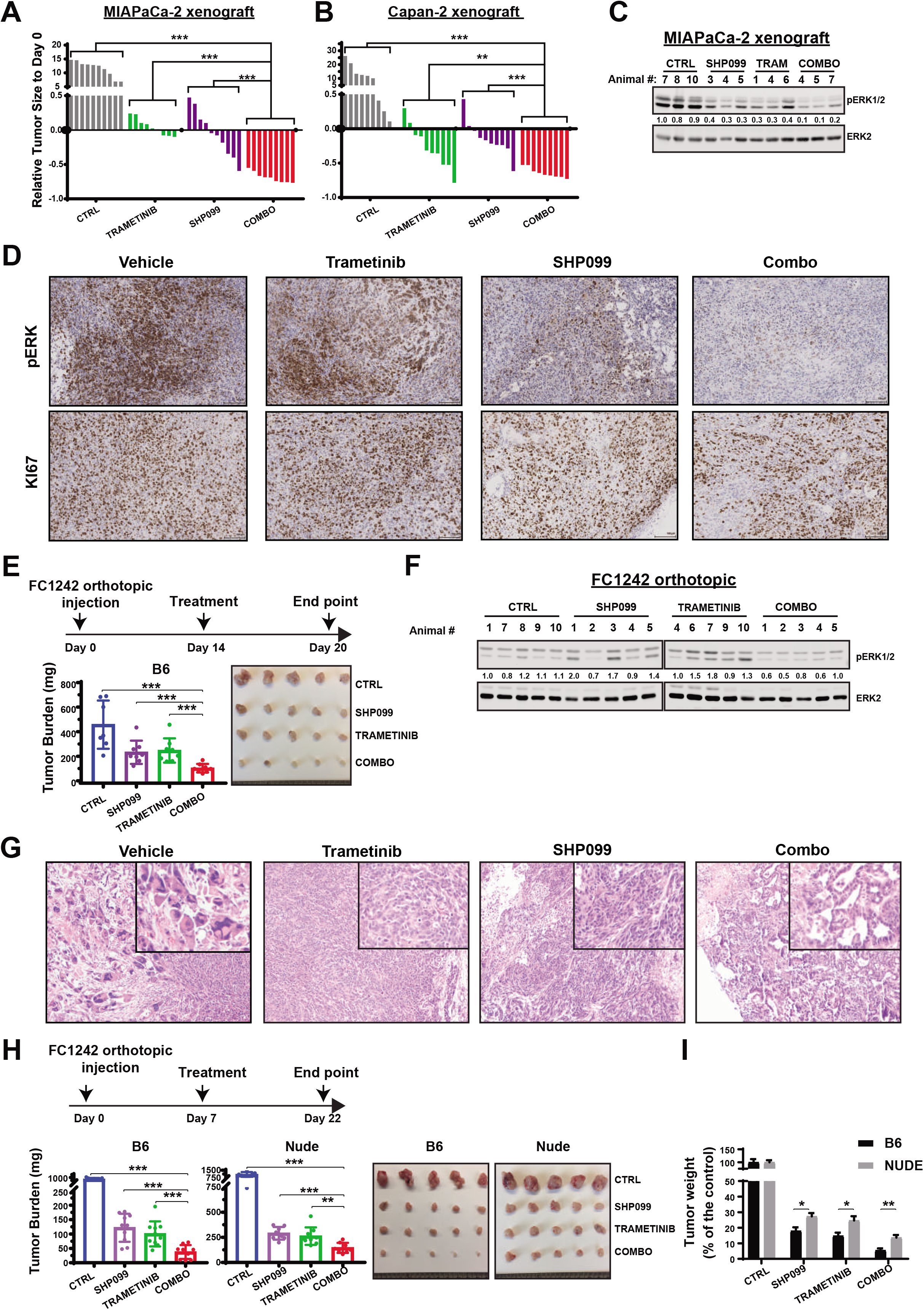
Combined MEK/SHP2 inhibition is efficacious in PDAC models *in vivo*. **A-B**, Response of MiaPaCa-2 (A) and Capan-2 (B) subcutaneous xenografts to treatment with SHP099 (75 mg per kg body weight, daily), trametinib (0.25 mg per kg body weight, daily) or both drugs (trametinib 0.25 mg/kg, daily; SHP099 75 mg/kg every other day). Waterfall plot shows response of each tumor after 3 weeks of treatment; *n* = 8-10 mice per group. (\*\**P*< 0.005; *** *P*< 0.0005, two-tailed Mann Whitney test). **C-D**, Effects on p-ERK levels in MiaPaCa-2 tumors following 3 weeks of drug treatment, as shown by p-ERK immunoblotting (C) and immunohistochemical analysis (D). **E**, Syngeneic mice with orthotopic injections of FC1242 (KPC) cells were treated with vehicle, SHP099 (75 mg per kg body weight, daily), trametinib (0.25 mg per kg body weight, daily) or both drugs (trametinib 0.25 mg/kg, daily; SHP099 75 mg/kg every other day), as depicted in the schematic, and tumor size was measured at day 14 (\*\*\**P*< 0.0005, one-tailed Mann Whitney test) **F**, Effects on p-ERK activity in FC1242-derived tumors from E. **G**, Representative haematoxylin and eosin (H&E) stains of drug-treated tumors from E. **H**, Sizes of tumors from syngeneic mice (C57BL/6J) and athymic nude (*nu/nu*) mice injected orthotopically with the same number of FC1242 (KPC) cells and treated with SHP099 (75 mg per kg body weight, daily), trametinib (1.0 mg per kg body weight, daily) or both drugs (trametinib 1.0 mg/kg + SHP099 75 mg/kg every other day), as depicted in the schematic, measured at day 22 (\*\**P*< 0.005, \*\*\**P*< 0.0005, one-tailed Mann Whitney test). **I**, Note larger effects seen in immune-competent mice (\**P*< 0.05, \*\**P*< 0.001; two-sided *t* test). Numbers under blots indicate relative intensities, compared with untreated controls, quantified by LICOR.

We also tested the drug combination in syngeneic mice injected orthotopically with a cell line (FC1242) (26) derived from induced LSL-Kras^G12D^*Trp53*^R172H^ (KPC) mice (27) (Fig. 3E). Tumors were allowed to grow for 14 days post-cell line injection, then mice were treated with the above single agent or combination regimen for 5 days and sacrificed. Again, mice in the combination arm had substantial tumor regressions, compared with trametinib-treated mice (Fig. 3E). Although the combination was clearly superior, SHP099 again had significant effects in this GEMM, even though KRAS(G12D) has significantly less residual GTPase activity than KRAS (G12C) (17). Immunoblot analysis again revealed more p-ERK inhibition in combination-treated tumors than in those treated with trametinib or SHP099 alone (Fig. 3F). Notably, the residual tumor mass in GEMMs treated with the drug combination exhibited significant exocrine cell differentiation, compared with those treated with vehicle or either single agent (Fig. 3G).

SHP2 is also implicated in signaling from multiple immune cell receptors, including checkpoint receptors such as PD-1 (28), so the anti-tumor effects of SHP099 alone or in combination could reflect drug action in the tumor microenvironment rather than, or in addition to, the tumor cells themselves. In exploratory studies to evaluate the effects of the immune system on the anti-tumor response to the trametinib/SHP099 combination, we compared the effects of drug treatment on orthotopic FC1242 tumors established in C57BL/6J and athymic nude (*nu/nu*) mice, respectively. Wild-type mice again exhibited significant tumor regressions in both single agent arms and a more potent anti-tumor effect in the combination arm (Fig. 3H). Immune-compromised hosts showed qualitatively similar responses to all treatment arms, but these responses were significantly lower that in immune-competent animals (Fig. 3I). Hence, an intact immune system is required for optimal tumor regression in response to all of these agents alone or in combination.

### SHP099/MEK-I combination is also effective in TNBC and serous ovarian cancer models

Genetic (29,30) and functional genomic (31,32) analyses have revealed striking similarities between TNBC and high-grade serous ovarian cancer (HGSC). Both of these malignancies typically express WT RAS, and in some TNBC models, MEK inhibition results in RTK upregulation and adaptive resistance (8). To explore the potential generality of combination MEK/SHP2 inhibition as a therapeutic strategy (and the utility of this combination in adaptive resistance to MEK-I in WT RAS-expressing cells), we tested SHP099 in combination with MEK-I in models of TNBC and HGSC.

Similar to its effects in *KRAS* mutant cells, MEK-I treatment increased the expression of various and variable RTKs and RTK ligand genes in TNBC and HGSC lines (Fig. S5A and B). Next, we tested these agents alone or in combination on a panel of TNBC and HGSC cell lines. SHP099 (10M) alone had little effect on cell number or colony formation (Fig. 4A-C). The MEK-I UO126 had variable effects, often (but not always) causing reduced cell proliferation compared with controls, but residual resistant cell populations were seen in almost all cell lines. The SHP099/MEK-I combination showed increased efficacy, with effects ranging from additive to synergistic (Fig. 4A and B). As in the *KRAS*-mutant cell models (above), combination treatment (for 48h) slowed cell cycle progression and enhanced cell death (Supplementary Fig. S5C and D).

**Figure 4.**
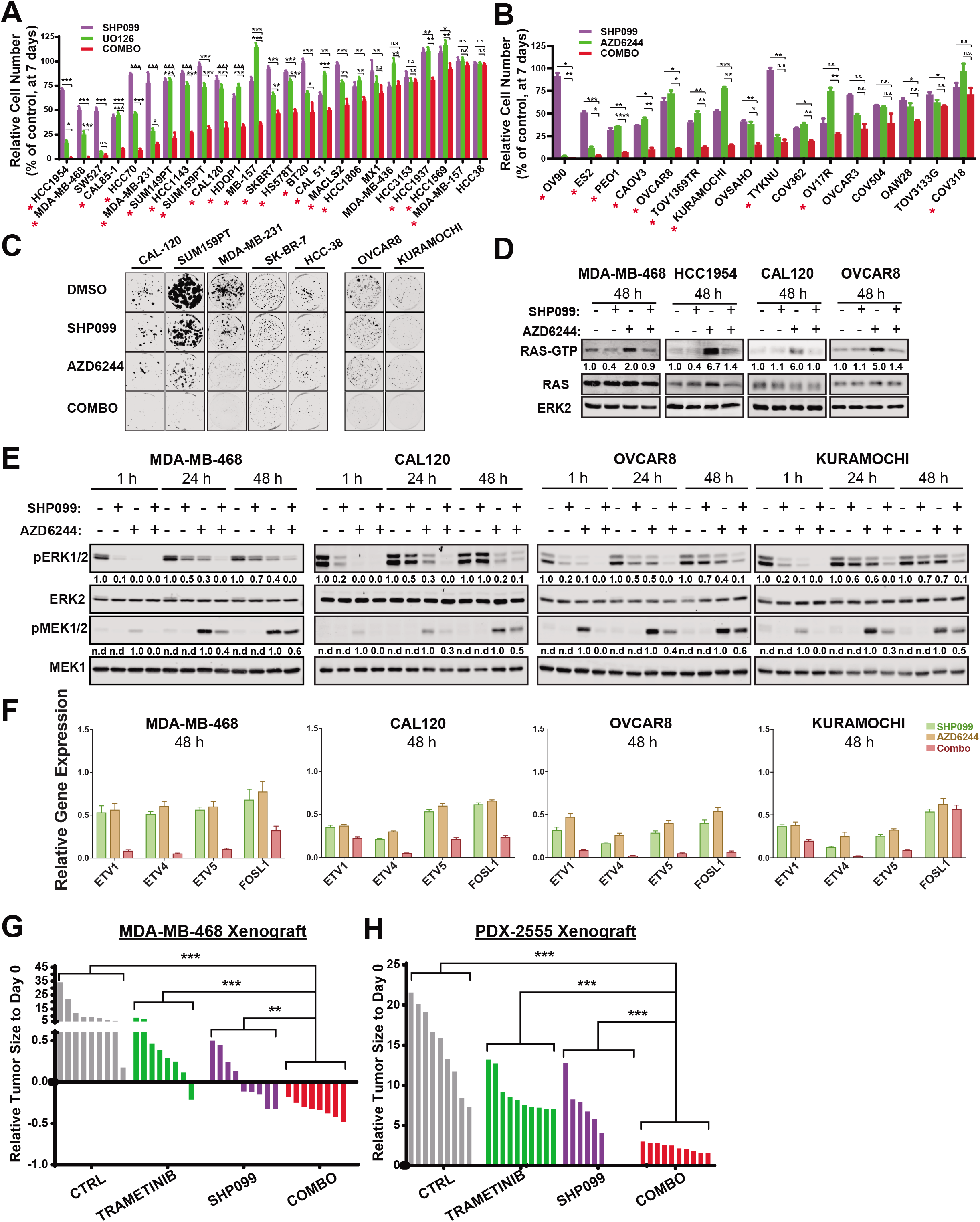
Combined MEK/SHP2 inhibition is also effective in TNBC and HGSC models *in vitro* and *in vivo*. **A-C**, TNBC (A) and HGSC (B) cell lines were treated with DMSO, SHP099, MEK-I (U0126 or AZD6244), or both (COMBO). PrestoBlue (A) and proliferation (B) assays were performed at one week. Colony formation (C) assay was performed at two weeks. Representative results from a minimum of three biological replicates are shown per condition: SHP099 10 μM, U0126 (10 μM) or AZD6244 (1 μM) as indicated, Combo=SHP099 10 μM + UO126 (10 μM)/AZD6244 1 μM (\**P*< 0.05, \*\**P*< 0.001, \*\*\**P*< 0.0001, two-sided *t* test). Red asterisk indicates synergistic interaction between the two drugs by BLISS independent analysis. **D**, GST-RBD pull down assay from TNBC and HGSC cell line lysates treated with DMSO, SHP099 10 μM, AZD6244 1 μM, or both for 48 h. Image is representative of at least two independent experiments. **E**, Immunoblots of lysates from TNBC and HSGS cell lines, treated as indicated. Image is representative of three independent experiments. **F**, ERK-dependent gene expression (*ETV1,4, 5* and *FOSL-1*), assessed by qRT-PCR, in TNBC and HGSC lines treated for 48h with the indicated drugs. **G-H**, MDA-MB-468 (G) and PDX-2555 (B) mammary fat pad xenografts following treatment with SHP099 (75 mg per kg body weight, daily), trametinib (0.25 mg per kg body weight, daily) or both drugs (trametinib 0.25 mg/kg, daily; SHP099 75 mg/kg every other day). Waterfall representations of tumor responses after 3 weeks of treatment are shown; *n* = 8-10 mice per group. (\*\**P* < 0.005, \*\*\**P*< 0.0005, two-tailed Mann Whitney test). Numbers under blots indicate relative intensities, compared with untreated controls, quantified by LICOR.

To examine the biochemical effects of these agents, we analyzed RAS/ERK pathway activation. After 48h of single-agent treatment, SHP099 had little or no effect on RAS activation in any of the models tested (Fig. 4D). After MEK-I treatment, however, RAS was hyper-activated to varying degrees in MDA-MB-468, HCC1954, CAL-120, and OVCAR-8 cells. In SHP099/MEK-I-treated cells, RAS-GTP decreased to the levels in randomly growing cells. These findings indicate that RAS activation is largely SHP2-independent under normal serum growth conditions, but is essential for the increase in RAS-GTP driven by the adaptive (RTK-driven) program evoked by MEK-I treatment. As expected, AZD6244 treatment for 48h also increased MEK1/2 phosphorylation, and consistent with its effects on RAS, SHP099 suppressed p-MEK (Fig. 4D). Similarly, SHP2 inhibition suppressed the adaptive increase in in p-ERK driven by 48h of MEK-I treatment (Fig. 4E), and the SHP099/MEK-I combination markedly suppressed ERK-dependent gene expression (Fig. 4F). Hence, our results suggest that SHP2 mediates RAS and ERK reactivation downstream of a diverse set of RTKs activated upon MEK inhibition in multiple cell systems, including those with mutant and WT RAS.

Finally, we treated mice bearing mammary fat pad xenografts derived from MDA-MB-468 or an extremely aggressive HGSC patient-derived xenograft (PDX) with SHP099, trametinib or the drug combination. Trametinib or SHP099 alone did not produce consistent regressions of MDAMB-468 xenografts, and the ovarian PDX was highly resistant to either agent. However, the SHP099/trametinib combination caused substantial regression of MDA-MB-468 xenografts and markedly inhibited the growth of the HGSC PDX (Fig. 4G). Thus, SHP2 inhibition suppresses reactivation of ERK1/2 downstream of different RTKs, which leads to tumor regression *in vivo*, suggesting that this combination approach might be broadly useful against acquired resistance to MEK inhibitors in cancer.

## DISCUSSION

Tumors evade targeted cancer therapies via an extensive repertoire of resistance mechanisms. One common theme involves activation of RTKs by upregulating their expression and/or the expression of their ligands, which leads to reactivation of the inhibited pathway (5-9,22,33). Consistent with these findings, multiple, distinct sets of RTKs/RTK ligands were activated in response to MEK-I treatment in the cancer cell models that we tested. The heterogeneity of this adaptive response renders unfeasible combination therapies with MEK-Is and RTK inhibitors. However, targeting a common downstream component of RTK signaling in combination with MEK-Is might yield substantial efficacy. SHP2 has long been known to signal downstream of normal RTKs (10,11), and cancer cells dependent on RTK activity are susceptible to SHP099 monotherapy (12,13). We find here that combined inhibition of MEK and SHP2 blocks cell proliferation and promotes tumor shrinkage in tumors with increased ERK pathway activation, including those that typically have WT RAS (TNBC, HGSC), as well as those driven by mutant KRAS (PDAC, NSCLC).

A recent paper suggested that SHP099 has off-target effects, at least in some cell contexts (34), but multiple lines of evidence indicate that SHP099 is “on-target” in our experiments. First, we observe the expected biochemical effects of this drug on RAS/ERK pathway activation in the multiple cell types tested. More compellingly, two different drug-resistant mutants of *PTPN11* (*PTPN11*^P491Q^ and *PTPN11*^*T253M/Q257L*^) rescue the effects of SHP099 in PDAC and NSCLC lines, respectively. Finally, the effects of a *PTPN11* shRNA phenocopy those of SHP099 biologically and biochemically. Similarly, the MEK-inhibitors employed here are highly validated, which strongly suggests that the effects we have observed here reflect dual SHP2/MEK inhibition.

We expected SHP099 to block the adaptive increase in *normal* RAS activation that accompanies increased RTK signaling in MEK-I-treated cells. Yet SHP099 also decreased *mutant* KRAS activation, as seen most clearly in MiaPaCA-2 cells, which only express KRAS(G12C), yet show clearly decreased KRAS activation following SHP099-treatment. KRAS(G12C) retains significant intrinsic GTPase activity *in vitro* (17), and this residual activity is significant *in vivo*, as exemplified by covalent RAS inhibitors that target KRAS(G12C)-GDP (35). Hence, even though KRAS(G12C) is largely refractory to RAS-GAPs, significant conversion to RAS-GDP must occur in cells via this intrinsic GTPase activity, and ongoing GDP/GTP exchange is required to maintain steady state levels of KRAS(G12C)-GTP. Other RAS mutants (with the exception of those affecting Q61) also retain some intrinsic GTPase activity, although much less than does KRAS (G12C). SHP099 also led to decreased RAS-GTP levels in cells expressing other KRAS mutants, yet whether that decrease also indicates an ongoing exchange requirement for these mutants, or instead reflects unloading of the remaining normal RAS proteins in those cells (36,37), remains unclear. Recently, Nichols *et al.* (38) reported variable effects of a new allosteric SHP2 inhibitor on mutant KRAS, although the chemical matter was not reported and its specificity was not established via drug-resistant mutations.

Regardless, our findings provide new insight into SHP2 action. Genetic and biochemical analyses firmly establish that SHP2 acts upstream of RAS (10,11), but whether it promotes exchange or inhibits GAP activity, or both, has been controversial. Early work showed that SHP2, via its C-terminal tyrosine phosphorylation sites, can recruit GRB2/SOS (39,40). A subsequent study reported diminished RAS exchange activity in lysates from cells expressing a mutant of GAB1 that cannot bind SHP2 in place of WT GAB1 (41). However, multiple other reports suggest that SHP2 antagonizes RAS-GAP activity by dephosphorylating its binding sites on RTKs or on SHP2-binding scaffolding adapters (42-44). Genetic evidence from studies of *Drosophila* also argue strongly for actions of the SHP2 ortholog, CSW, on the GAP binding sites in TORSO (45). Our findings, and those of Nichols *et al.* (38), show clearly that SHP2 acts upstream of SOS, although we cannot exclude additional effects on RAS-GAP.

Although SHP099 has little effect on 2D proliferation or colony assays, it significantly affected some xenografts and the KPC GEMM. Several potential, non-mutually exclusive explanations exist for this apparent discrepancy. First, tumors exist in a hypoxic, nutrient-challenged, and potentially growth-factor deficient microenvironment; under such conditions, SHP2 function might be essential. Second, SHP2 might affect stromal support functions (e.g., growth factor production by stromal fibroblasts). Third, SHP2 might affect the anti-tumor immune response. In this regard, our finding that SHP099 (and trametinib and combo) efficacy is enhanced in immune-competent, compared with *nu/nu*, mice indicates a significant role for T cells in opposing the KPC tumor growth.

Our results comport with, and extend, two previous studies of the effects of SHP2 modulation on other ERK pathway inhibitors. Prahallad *et al.* (14) found that SHP2 depletion (via *PTPN11* shRNA or deletion) blocked adaptive resistance to the BRAF^V600E^ inhibitor vemurafenib. They claimed that an SHP2 catalytic domain inhibitor (GS493) had similar effects, but that agent also inhibits tyrosine kinases (46). During the preparation of this manuscript, Dardaei *et al.* (47) found that SHP2 depletion or SHP099 treatment could resensitize crizotinib- and ceritinib-resistant *ALK*-mutant NSCLC cell lines. Together with our studies, these results suggest that SHP2 inhibition might be a broadly applicable strategy to prevent or overcome adaptive resistance to kinase inhibition in a wide array of malignancies.

## METHODS

### Cell lines and reagents

All cell lines were grown at in 5% CO_2_ at 37C° under the conditions described by the vendor or the source laboratory; details are available from CF and KHT upon request. Cells were tested routinely (every 3 months) for mycoplasma contamination by PCR (48), and genotyped by STR analysis at IDEXX Bioresearch.

HCC1954, MDA-MB-468, SW527, CAL85-1, HCC70, MDA-MB-231,SUM149PT, HCC1143, SUM159PT, CAL120, HDQP1, MB-157, SKBR7, HS578T, BT20, CAL51, MACSL2, HCC1806, MX1, MDA-MB-436, HCC3153, HCC1937, HCC1569, MDA-MB-157 and HCC38 are from laboratory stocks, obtained as described previously (31). AsPC1, CFPAC-1, DAN-G, ES-2 Hs766T, HuPT3, Panc02.03, Panc02.13, Panc04.03, Panc08.13, Panc10.05, PSN-1, and SU.86.86 were purchased from ATCC. Capan-2, CFPAC-1, HPAC, HPAF-II, HuPT4, MIAPaCa-2, Panc03.27, PATU8902, and SW1990 cells were obtained from Dr. Alec Kimmelman (NYU School of Medicine). Caov3 cells were obtained from Dr. Douglas Levine (NYU School of Medicine). KURAMOCHI, OAW28, OVSAHO, OV17R and TYKnu cells were obtained from Dr. Gottfried Konecny (UCLA). COV318, COV362, COV504, OV90, OVCAR3, OVCAR8, PEO1, TOV1369TR and TOV3133G cells were obtained from Dr. Robert Rottapel (Princess Margaret Cancer Center). KPA, KPC, CALU-1, H23 and H358 lung cancer cell lines were obtained from Dr. Kwok-Kin Wong (NYU School of Medicine).

The NYU 16, 53 and 59 primary low passage human pancreatic cancer cell lines were generated according to Institutional Review Board (IRB) guidelines from invasive pancreatic adenocarcinoma samples from patients who underwent surgical resection at University of Michigan Hospital and New York University. In brief, cell lines were generated by xenotransplantation of human PDA into immune deficient mice; tumor cells were isolated using Magnetic Cell Isolation followed by MACS technology (Miltenyi Biotech). Cells were then plated and subcloned in RPMI 1640 medium, supplemented with 10% heat-inactivated FCS (Gibco), 2 mm l-glutamine, 100 units/ml penicillin, 100 μg/ml streptomycin (Invitrogen) in 95% air and 5% CO2 at 37 °C. Cultures were subsequentially purified of mouse cell stroma by using a mouse cell depletion kit (Miltenyi Biotech) and purity confirmed by FACS analysis.

SHP099 (HY-100388A) was purchased from MedChemExpress. Selumetinib-AZD6244 (S1008), UO126 (S1102) and trametinib (GSK1120212-S2673) were purchased from Selleckchem.

### Plasmids, retro and lentiviral production

A human SHP2 cDNA was cloned into pMSCV-IRES-GFP or pCW57.1 (pCW57.1 was a gift from David Root, Addgene plasmid #41393). Mutations were introduced by using the QuikChange II site-directed mutagenesis kit (Agilent Technologies). The IPTG inducible lentiviral shRNA plasmid vector pLKO.1-901 was obtained from Dr. Jason Moffat (University of Toronto), and shRNA against human *PTPN11* (shSHP2) (5'CGCTAAGAGAACTTAAACTTT 3') and a control shRNA against GFP (5' TGCCCGACAACCACTACCTGA 3') were introduced into this vector. The *SOS1* B1 (catalytic domain of SOS fused with a C-terminal CAAX T7 tag) construct was a gift from Dr. Dafna Bar-Sagi (NYU School of Medicine).

Viruses were produced by co-transfecting HEK293T cells with lentiviral or retroviral constructs and packaging vectors (pVSV-G + pvPac for retroviruses; pVSV-G + dR8.91 for lentiviruses). Forty-eight (48) hours later, culture media were passed through a 0.45 mm filter, and viral supernatants, supplemented with 8 g/ml of polybrene (Sigma), were used to infect 70% confluent cells in six-well dishes for 16h at 37 °C. Stable pools were selected either by using the appropriate antibiotic or by fluorescence activated cell sorting (FACS) for EGFP.

### PrestoBlue Assay

Cancer cells were seeded in 96-well tissue culture plates (500-3,000 cells/well). Following incubation with DMSO, 1 μM AZD6244, 10 μM SHP099 or both drugs, cell viability was assayed at 0, 1, 3 and 7 d (n =3) using the PrestoBlue cytotoxicity assay (Thermo Fisher), according to the manufacturer’s protocol. Media (including drugs) were refreshed every 48 h. Briefly, 10 μL of PrestoBlue reagent were added to each well, and after 2h, fluorescence was measured on a multi-plate reader, using excitation wavelength 530 nm/emission wavelength 590 nm. Data were corrected for PrestoBlue background fluorescence in media alone. All data represent at least three biological independent experiments.

### Clonogenic survival assay

Cells (100 to 500) were seeded in six-well plates one day before treatment with DMSO, 1 μM AZD6244, 10 μM SH099 or the drug combination, allowed to grow until they formed colonies (7-14 days), rinsed twice with PBS to remove floating cells, fixed in 4% formaldehyde in PBS (v/v) for 10-15 minutes, and stained in 0.1% crystal violet/10% ethanol for 20 minutes. Staining solution was aspirated, and colonies were washed with water 3x, air-dried and visualized with the Odyssey Imaging System (LICOR). Results were quantified by using the ImageJ Colony Area PlugIn (49). At least three biological replicates were performed.

### RAS activity measurements

Cells (5 × 10^6^) in 15-cm dishes were treated with the indicated concentrations of compound(s) for the times indicated. One (1) mL of lysis buffer (25 mM Tris-HCl, pH 7.2, 150 mM NaCl, 5 mM MgCl_2_, 5% glycerol, 1% NP40) containing protease inhibitors was added for 15 minutes on ice, and lysates were scraped from the plate and centrifuged at 14,000 rpm for 15 minutes at 4°C. Clarified lysates (3 mg) were added to pre-washed GST-tagged RBD glutathione agarose beads (30 μL) for 1 hour at 4 °C under constant rocking. Beads were then centrifuged, washed once, and eluted in 30 μL of 2x SDS-PAGE sample buffer. Immunodetection of RAS proteins was carried out with KRAS antibody (F234: sc-30; Santa Cruz Biotechnology; 1:250), NRAS (F155: sc3-1; Santa Cruz Biotechnology; 1:250) or pan-RAS antibody (Ab-3; Calbiochem; 1:1,000), with ERK-2 (D2: sc-1647; Santa Cruz Biotechnology; 1:1000) immunstaining used as a loading control.

### Immunoblotting

Whole cell lysates were generated in modified radioimmunoprecipitation (RIPA) buffer (50mM Tris-HCl pH 8.0, 150mM NaCl, 2mM EDTA, 1% NP-40, and 0.1% SDS, without sodium deoxycholate), supplemented with protease (40μg/ml PMSF, 2μg/ml antipain, 2μg/ml pepstatin A, 20μg/ml leupeptin, and 20μg/ml aprotinin) and phosphatase (10mM NaF, 1mM Na_3_VO_4_, 10mM β-glycerophosphate, and 10mM sodium pyrophosphate) inhibitors. After removal of debris, samples were quantified with the DC Protein Assay Kit (Bio-Rad). Total lysate protein was resolved by standard SDS-PAGE, and transferred in 1X transfer buffer and 15% methanol. Membranes were incubated with their respective primary and secondary antibodies labeled with IRDye (680nm and 800nm) and visualized using the LICOR. Antibodies against phospho-p42/44 MAPK (rabbit polyclonal; #9101; 1:1000), phospho-MEK 1/2 (rabbit polyclonal; #9121; 1:1000), MEK 1 (61B12; mouse monoclonal; # 2352; 1:1000) and T7-tag (D9E1X; rabbit monoclonal; # 13246) were obtained from Cell Signaling. Rabbit polyclonal antibodies against SHP2 (sc-280; 1:1000) and mouse monoclonal ERK-2 (D2: sc-1647; 1:1000) were purchased from Santa Cruz Biotechnology. Mouse monoclonal Ab anti SOS-1 (# MA5-17234) was purchased from Invitrogen.

### Cell Cycle Analysis

Cells were fixed in cold 70% ethanol overnight, washed with PBS, then stained with buffer containing 20uL 7AAD (BD) and RNase (final concentration 0.5ug/mL) for 60 minutes in room temperature. Stained cells were analyzed by flow cytometry on LSR II machine (BD). Data was then analyzed by ModFit LT software (Versity Software House) to determine fraction of cells in each cell cycle stage.

### Annexin V Analysis

Cells were stained with PE Annexin V Apoptosis Detection Kit I according to manufacturer’s protocol (BD). Stained cells were analyzed by flow cytometry on LSR II machine (BD), and data were analyzed by FlowJo software (BD).

### Xenografts

All animal experiments were approved by the NYU Langone Institutional Animal Care and Use Committee (IACUC). MiaPaCa-2 and Capan-2 xenografts were established by subcutaneous injection of 5 × 10^6^ cells in 50% Matrigel (Corning) into the right flank of nude mice (nu/nu, #088 Charles River) when animals were 8 to 10 weeks of age. MDA-MB-468 xenografts were established by injecting 5 × 10^6^ cells in 50% Matrigel into the right lower mammary pad. Ovarian PDXes were established by injecting 5 × 10^5^ cells in 50% Matrigel into the right lower mammary pad of NSG mice (Jackson Lab) when animals were 6 to 8 weeks of age.

Each treatment group contained 8-10 mice. When tumors reached 100-500 mm^3^, as measured by calipers (size=length*width^2^*0.5), mice were randomized to four groups (10 mice/group) for each model, and treated with: (i) vehicle, (ii) SHP099, (iii) trametinib, or (iv) SHP099/trametinib. Investigators were not blinded to group allocation. The following oral gavage dosing regimens were employed: SHP099 75mg/kg QD, trametinib 0.25mg/kg QD, and trametinib 0.25mg/kg QD, SHP099 75mg/kg QOD. SHP099 was resuspended in 0.6% methylcellulose, 0.5% Tween80 in 0.9% saline. Trametinib was dissolved in DMSO before adding to the carrier. Caliper and weight measurements were performed every other day and continued until termination of the experiments.

### Syngeneic orthotopic pancreatic cancer models

FC1242 cells were generated in the Tuveson lab (Cold Spring Harbor Laboratory) from a pancreas tumors in a LSL-*Kras*^G12D^; LSL-*Trp53*^R172H^; Pdx1-Cre (KPC) mouse on C57BL/6 background, as described (27). Cells (1 x 10^5^) were suspended in Matrigel, implanted into the pancreata of 6-8 week-old syngeneic male mice, and allowed to establish for 1-2 weeks before beginning treatment. Vehicle, trametinib (0.25mg/kg QD), SHP099 (75mg/kg QD), or trametinib 0.25 mg/Kg QD, SHP099 75 mg/kg QOD was administered for five days, and mice were sacrificed.

### qRT-PCR

Total cellular RNA extraction was isolated by using the Qiagen RNeasy kit. cDNA was generated by using the SuperScript IV First Strand Synthesis System (Invitrogen) for RT-PCR. qRT-PCR was performed with Fast SYBR™ Green Master Mix (Applied Biosystems), following the manufacturer’s protocol, in 384-well format in C1000 Touch Thermal Cycler (Biorad). Subsequent differential gene expression analysis was performed with CFX Manager (Biorad) and normalized to GAPDH expression. Primers used are listed in Supplementary Table 1.

### Bliss Analysis

Potential syngergistic effects of SHP099 and AZD6244 were determined by Bliss analysis as *Y*_*ab,P*_ = *Y*_*a*_ + *Y*_*b*_ – *Y*_*a*_*Y*_*b*_, where *Y*_*a*_ stands for percentage inhibition of drug *a* and *Y*_*b*_ stands for percentage inhibition of drug *b* (50).

## Abbreviations

HGSC, high-grade serous ovarian cancer; MAPK, mitogen-activated protein kinase; MEK-Is, MEK-inhibitors; NSCLC, non-small cell lung cancer, PDAC, pancreatic ductal adenocarcinoma; TNBC, triple negative breast cancer

## Acknowledgement

We thank Drs. Alec Kimmelman, Douglas A. Levine, Gottfried E. Konecny, Robert Rottapel, and Kwok-Kin Wong for sharing cell lines, Drs. Dafna Bar-Sagi and David Root for plasmids, and the PCC Experimental Pathology and Precision Immunology shared resources for technical support. We also thank Dr. Toshiyuki Araki for advice and discussion on this project. This work was supported by R01CA49152 and P30CA016087 to B.G.N., and R01CA131045 to D.M.S‥

